# *Switching Gō-Martini* for Investigating Protein Conformational Transitions and Associated Protein-Lipid Interactions

**DOI:** 10.1101/2023.08.21.554122

**Authors:** Song Yang, Chen Song

## Abstract

Proteins are dynamic biomolecules that can transform between different conformational states when exerting physiological functions, which is difficult to simulate by using all-atom methods. Coarse-grained Gō-like models are widely-used to investigate large-scale conformational transitions, which usually adopt implicit solvent models and therefore cannot explicitly capture the interaction between proteins and surrounding molecules, such as water and lipid molecules. Here, we present a new method, named *Switching Gō-Martini*, to simulate large-scale protein conformational transitions between different states, based on the switching Gō method and the coarse-grained Martini 3 force field. The method is straight-forward and efficient, as demonstrated by the benchmarking applications for multiple protein systems, including glutamine binding protein (GlnBP), adenylate kinase (AdK), and *β*_2_-adrenergic receptor (β2AR). Moreover, by employing the *Switching Gō-Martini* method, we can not only unveil the conformational transition from the E2Pi-PL state to E1 state of the Type 4 P-type ATPase (P4-ATPase) flippase ATP8A1-CDC50, but also provide insights into the intricate details of lipid transport.

## 1 Introduction

Proteins perform a vast variety of fundamental functions in living systems, including catalyzing enzymatic reactions, signal transduction, and transporting molecules. During most of these processes, proteins need to change their structures to dynamically adapt to their functions. For example, enzymes enclose the catalytic pockets to catalyze the reactions after binding to the substrates. ^1,2^ Mechanically activated ion channels can open the pores to allow ions passing through the membrane when sensing the increase of surface tension. ^3^ GPCRs sense the binding of small molecules followed by the conformational changes of the sixth transmembrane helix to induce the G protein binding and signal transduction. ^4^ Thus, proteins in living organisms need to be dynamic to undergo conformational transitions to exert their functions.

The conformational changes of proteins can be monitored by super-resolution experimental methods and computational simulation methods. The structural biology methods, including X-ray crystallography, nuclear magnetic resonance (NMR) and cryo-electron microscopy (cryo-EM), can primarily capture the most populated, ensemble-averaged structures, although efforts have been made to uncover the high-energy, low-populated conformations during protein dynamics. ^5–7^ Single-molecule technologies have also been used to characterize the conformational transitions of proteins, such as single-molecule Förster resonance energy transfer (smFRET) and single-molecule force spectroscopy (SMFS). ^8^ However, these methods merely capture some low-dimension characteristics of proteins and cannot describe the full conformational changes of proteins at atomic resolution. To alleviate the detail loss in wet experiments, molecular dynamics (MD) simulation has been the most widely-used computational method for revealing the transition paths of proteins. All-atom MD simulations are often limited to the microsecond timescale, so a variety of enhanced sampling methods are developed to accelerate the exploration of protein transitions, such as replica exchange, ^9^ metadynamics, ^10^ and umbrella sampling. ^11^ In addition, the Markov state model is often utilized to sample the transition paths of protein conformational changes. ^12^ Nonetheless, it is still challenging to sample the large-scale conformational transitions of proteins, and the expensive computational cost does not allow high-throughput applications.

A potential solution is the use of coarse-grained (CG) force fields, which reduces the computational cost by uniting groups of atoms into effective interaction sites, resulting in a substantial computational speed-up. One of the most popular CG methods is the Gō-like model, which groups one residue or more to a pseudo atom with Lenard-Jones potential to maintain the second and higher structures. ^13^ These methods have obtained great success in simulating the conformational changes of glutamine-binding protein, ^14^ adenylate kinase, ^15,16^ molecular motor protein, ^17^ and so on. However, the lacking of an explicit solvent environment and biomembrane in these models may impede the system from accurately simulating the actual dynamic process of membrane proteins, especially when the interactions of proteins and lipids are crucial, which is often the case.

The Martini force field is one of the most widely-used CG force fields in simulating the interaction between proteins and lipids. ^18,19^ However, this force field was hardly used to simulate protein conformational changes because the previous versions utilized the elastic network potential, which hindered large conformational changes to occur unless heavily adapted. ^20^ Recently, the Gō-like Martini model, which uses the Lennard-Jones potential network to replace the elastic network, showed very promising results in studying protein dynamics. ^21,22^ With the Martini 3 released, it was further demonstrated that the switching approach can be utilized to study small molecular switches. ^23^ Therefore, it appears that the combination of the Switching Gō-like model with the Martini 3 force field may carry a great potential in simulating conformational changes of multiple-state proteins, similarly to the previous Gō models, ^13,14^ but with explicit surrounding molecules.

In this study, we present that the switching approach for the Gō-like Martini model can indeed be used to simulate conformational changes of proteins and sample the transition path with high accuracy and fast speed. We utilized three well-tested proteins, including GlnBP, AdK, and β2AR, to benchmark the accuracy of our method. Then, we applied the method to investigate the lipid transport mechanism of the phosphatidylserine (PS) flippase ATP8A1-CDC50a. The process of lipid translocation from the exoplasmic leaflet to the cytoplasmic leaflet was successfully simulated, which was facilitated by conformational transitions of the protein. Our results demonstrate that, with the *Switching Gō-Martini* method, one can not merely efficiently sample the potential conformational transition paths of proteins, but also gain insights into the interactions between dynamic proteins and lipids.

## 2 Results

### 2.1 Conformational transitions of GlnBP, AdK and β2AR

The switching method is a straightforward way to transform the energy landscape of one state of protein to the other, which has been extensively utilized in previous studies. ^17,24–26^ The schematic of Fig. 1a illustrates the mechanism of the switching approach. While the protein oscillates in the energy well A with the energy landscape of State A, we manually switch the energy landscape of the protein to that of State B. After that, the protein structure of State A has high energy and becomes unstable. Gradually, the protein can change its conformation to fall into the energy well B. The details of the implementation can be found in the section of Materials and Methods. To validate the approach, we tested it in three well-studied benchmarking systems, including GlnBP, AdK, and β2AR.

**Figure 1:**
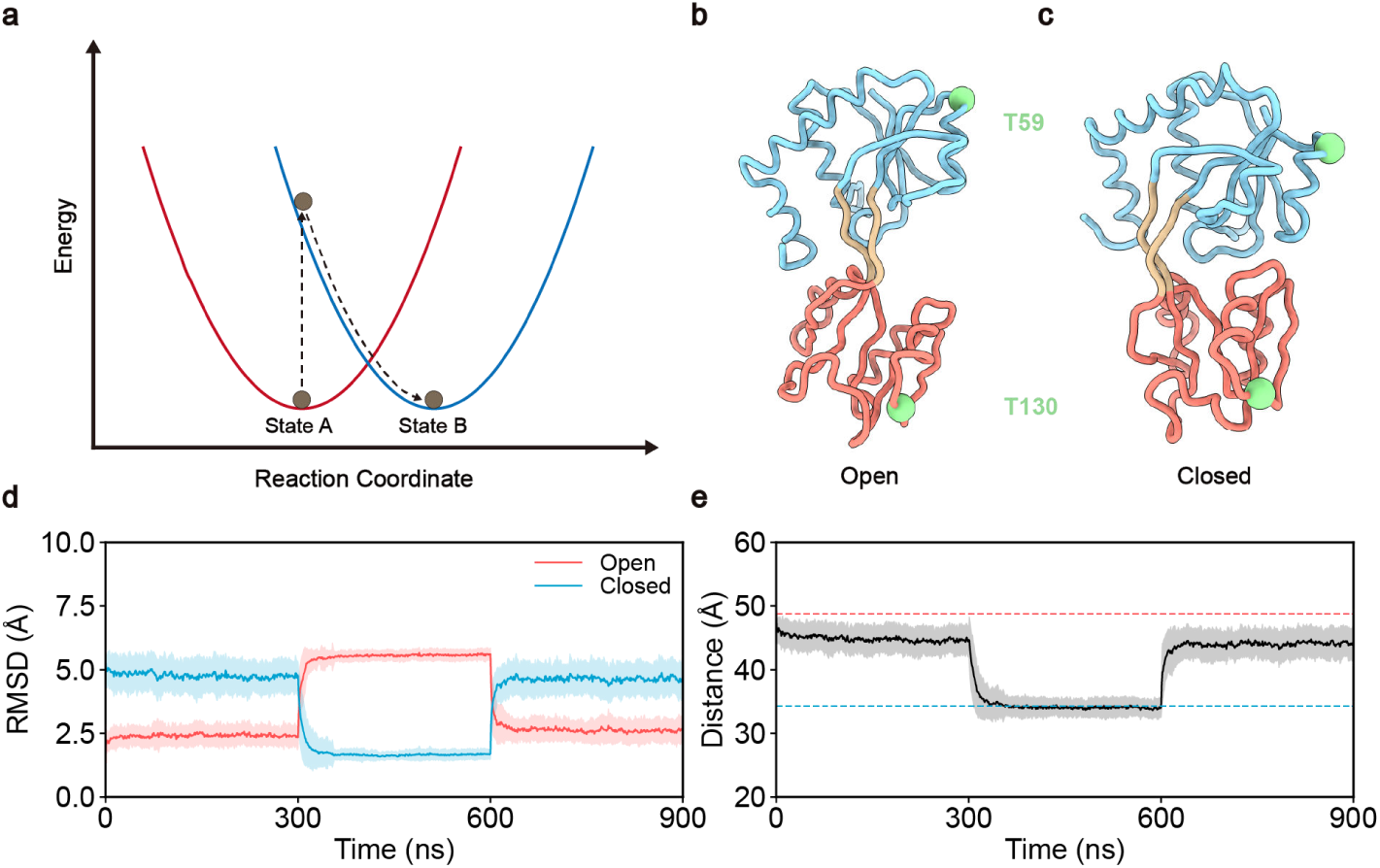
Switching Gō-Martini simulation for GlnBP. **a,** Schematic of the Switching Gō-Martini approach. The energy landscapes are simplified as quadratic curves (blue and red) and the protein state is drawn as a ball (brown). **b, c,** The coarse-grained structures of GlnBP in the open and closed state, respectively. **d,** The RMSD of GlnBP during the simulations, with respect to the open-(red) or closed-state (blue) structures. The solid lines correspond to the average values obtained from 50 independent simulation trajectories, while the shaded area represents the standard deviation. **e,** The distance between the backbone atoms of T59 and T130 during the Switching Gō-Martini simulations. The solid line represents the average value, similar to **d**, and the shaded area represents the standard deviation. The red and blue dashed lines represent the reference distance from the open- and closed-state structures.

#### 2.1.1 GlnBP

GlnBP is one of the periplasmic binding proteins, which assists L-glutamine uptake in bacteria. ^1^ The crystal structures of GlnBP were resolved with a ligand-bound closed state and a ligand-free open state (Fig. 1b, c). ^27,28^ In the holo state, the glutamine is surrounded by two global domains (large domain and small domain) connected by a linker. Both the experimental tests and simulation models have revealed the open–closed transition of GlnBP during the glutamine associating and disassociating process. ^29–31^

We used the Switching Gō-Martini approach to simulate 50 independent repeats of the conformational changes of the GlnBP from open to closed and back to open state in our system. The root-mean-square deviation (RMSD) of CG backbone (BB) atoms was used to show the conformational changes of GlnBP during our simulations. As shown in Fig. 1d, the results of 50 independent simulations showed that the conformation of GlnBP changed along the expected transition pathway when we switched on the open- or closed-energy landscapes of GlnBP. The opening extent of the substrate binding pocket was measured by the distance between the BB atoms of T59 (large domain) and T130 (small domain). As expected, the distance in the simulations decreased after the closed state was switched on and increased when the open state was switched on (Fig. 1e). These results demonstrate that the Switching Gō-Martini method can indeed be utilized to simulate the conformational transitions between two distinct states.

To further demonstrate the robustness of the approach, a continuous simulation comprising ten cycles of open↔closed transitions of GlnBP was performed. Throughout the simulation, both the RMSD and distance of T59-T130 changed with the open or closed state switched on or off (Fig. S1a, b). Moreover, the metrics associated with the open or closed state remained stable and consistent during the simulations. These results indicate that the Switching Gō-Martini method is robust in simulating repetitive conformational transitions between distinct states.

#### 2.1.2 AdK

AdK is a ubiquitous enzyme that catalyzes the reversible phosphoryl transfer of AMP and ATP into two ADP, which is important in cellular energy homeostasis. ^2^ AdK consists of the ATP-binding domain (LID), AMP-binding domain (NMP), and the remainder linking them named as Core domain. The crystal structures of the closed and the open states of AdK in Fig. 2a, b show that the LID and NMP keep closed when binding to ATP and AMP while exposing their binding pockets in the unbound state. ^32,33^ The conformational transition of AdK between the open and closed states is frustrated and in a stepwise manner, which has been widely studied by previous experiments and computational simulations. ^15,16,34^ There are two major kinetic pathways found, the NMP-closing pathway and the LID-closing pathway. The NMP-closing pathway is characterized by the intermediate structure in which the NMP keeps closed and LID open (Fig. 2c), while in the LID-closing pathway the NMP domain is open and LID closed (Fig. 2d).

**Figure 2:**
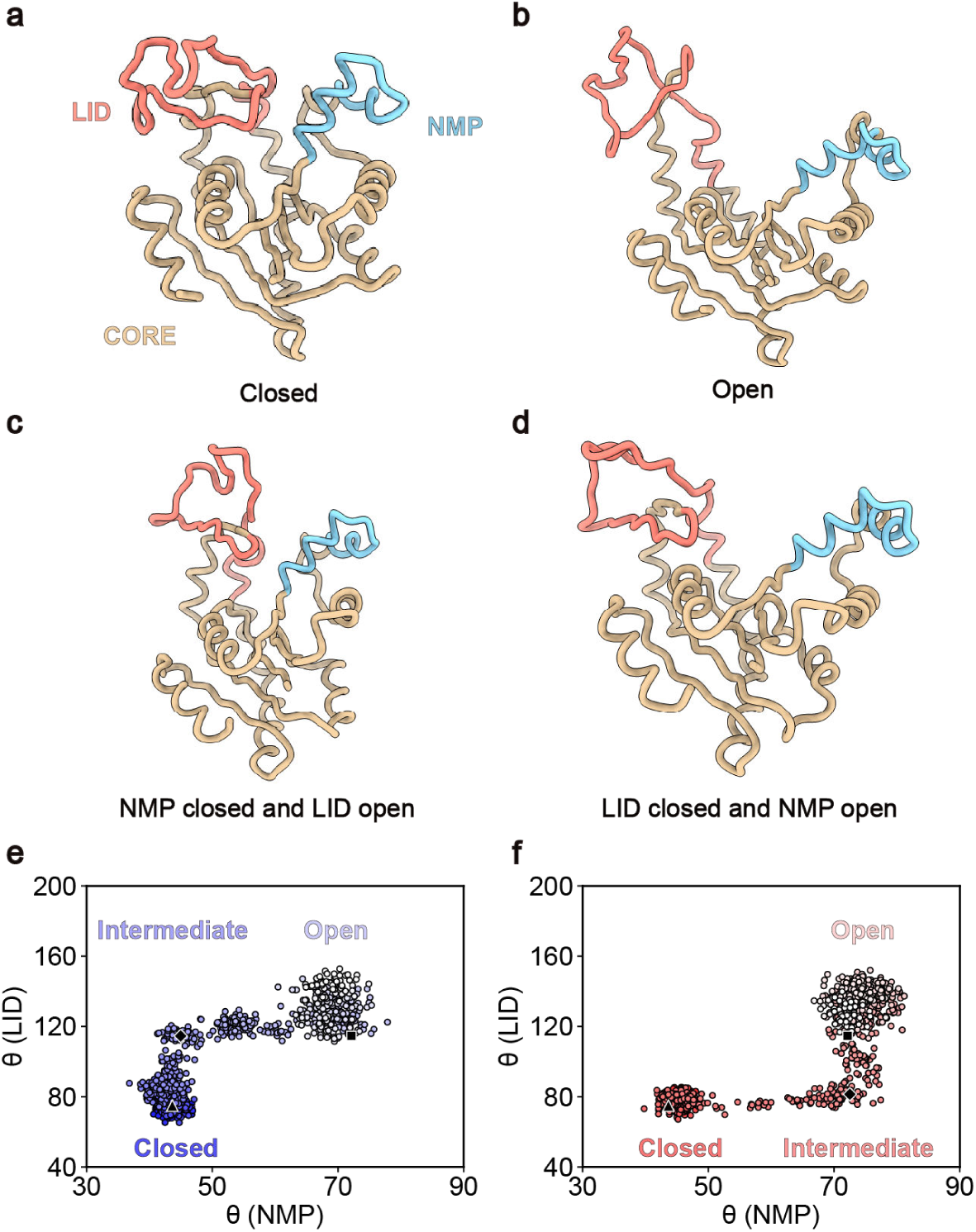
Switching Gō-Martini simulation for AdK. **a, b, c, d,** Representative structures of four major conformational states with NMP and LID open or closed. NMP domain (residues 31-60), LID domain (residues 127-164), and the remainder are colored sky blue, salmon, and tan, respectively. **e, f,** The two-dimension conformational transition paths (NMP-closing pathway, left; LID-closing pathway, right) are constructed with two angles *θ_NMP_* and *θ_LID_* as the reaction coordinates. The color of the circles decreases along with the simulation time. The black triangle, diamond, and square indicate the closed, intermediate, and open states of AdK.

We used the Switching Gō-Martini method to search the above pathways in 70 independent closed-to-open MD simulations, starting with the closed structure. The RMSD of AdK showed that conformational changes occurred when we switched on the open-state energy landscape (Fig. S2a). We used two angles *θ_NMP_* and *θ_LID_* as reaction coordinates to characterize the state of the ensembles during our simulations (Fig. 2e, f and Fig. S2b–f). The analysis of the conformational changes revealed that the LID domain and NMP domain underwent a gradual opening during the transition process. Both the NMP-closing pathway and LID-closing pathway were found in our Switching Gō-Martini simulations, and the NMP-closing pathway was dominant with a probability of 68.6%, compared to 10% along the LID-closing pathway (Fig. S2d, e). Our results for AdK’s NMP-closing pathway are consistent with the previous computational works by Lu et al., Li et al. and Stiller et al. ^15,35,36^ However, it is noteworthy that there are also papers reporting the LID-closing intermediate to be more stable. ^37,38^

#### 2.1.3 β2AR

The third benchmark is the deactivation of β2AR, a member of the transmembrane protein family of G protein-coupled receptors (GPCR). β2AR functions by binding to adrenaline and transducing the signal into the cell, playing important roles in muscle relaxation, vasodilation, and insulin secretion. ^4,39^ The activation and deactivation mechanisms of β2AR have been experimentally well studied, in which the snapshots of the inactive and active state β2AR structures (Fig. 3a, b) have been characterized by the X-ray method. ^40,40,41^ However, the detailed conformational transition pathway of the β2AR activating and deactivating process remained elusive. Therefore, micro- to milli-seconds all-atom simulations were conducted to reveal the activation and deactivation mechanism of β2AR, ^42,43^ which provided a solid basis for benchmarking Gō-like coarse-grained simulations.

**Figure 3:**
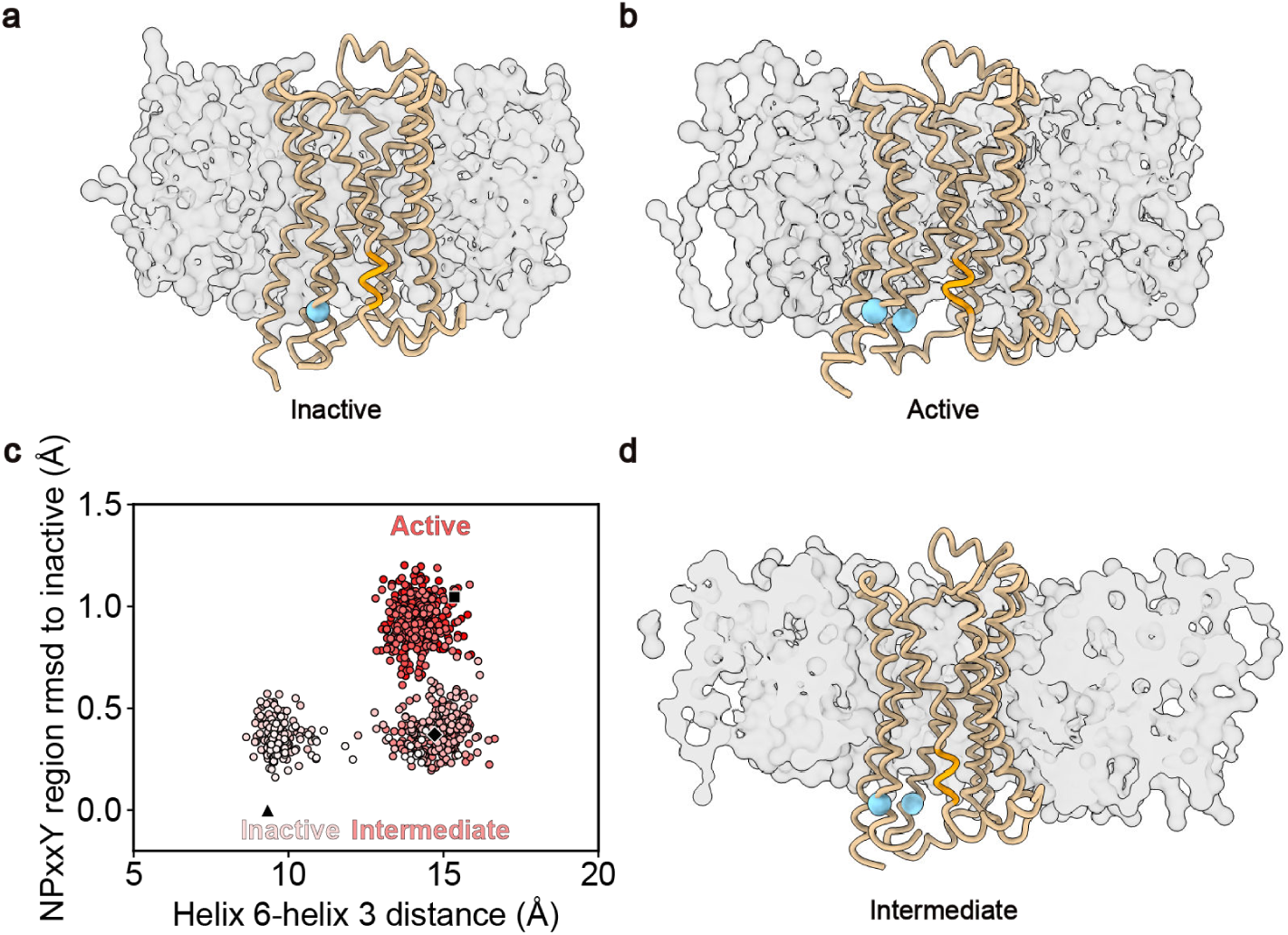
Switching Gō-Martini simulation for β2AR. **a, b, d,** Representative coarse-grained structures of β2AR in the inactive, active, and intermediate states embedded into the POPC bi-layer, respectively. The protein is shown as the tan ribbon, the NPxxY motif is colored orange, and the BB of R131 and L272 are colored sky-blue with sphere style. The POPC membrane is shown as gray surfaces. **c,** Two-dimension conformational transition pathway, as constructed with the helix 6–helix 3 distance and the RMSD of NPxxY motif with respect to the inactive state. The red color of the circles decreases along with the simulation time. The black square, diamond, and triangle are representative of the active, intermediate, and inactive states of β2AR.

We used the Switching Gō-Martini method to simulate the deactivation process of β2AR for 40 independent repeats and compared our results with those from previous works. The RMSD of BB atoms showed that the conformational transitions took place after we switched on the energy landscape of the inactive state of β2AR (Fig. S3a). To accurately analyze the conformational transition path, the distance between helix 6 and helix 3 (measured between residues R131 and L272) and the RMSD of the NPxxY motif in helix 7 with respect to the inactive structure were utilized as the reaction coordinates. Fig. 3c showed that the helix 6–helix 3 distance was 14.0 ± 0.6 Å in the active state and gradually decreased to 9.4 ± 0.5 Å which was within the range of the inactive state. The RMSD of the NPxxY motif changed and was finally close to the inactive structure. Unlike the conformational changes of AdK during the catalysis, β2AR mainly underwent a single pathway during the deactivation process, in which the RMSD of NPxxY firstly decreased and the intracellular end of helix 6 gradually shifted toward helix 3 (Fig. 3c, d and Fig. S3b, c). Only one rare pathway was found within our 40 simulations, in which the order of conformational transitions on the two reaction coordinates was reversed (Fig. S3d). All these observations were consistent with the unbiased all-atom MD simulations using the specialized supercomputers Anton. ^42^

#### 2.1.4 Benchmark summary

We utilized three well-tested examples including GlnBP, AdK, and β2AR to examine the Switching Gō-Martini method. The results show that our approach can not only simulate the conformational changes of protein between different states in the presence of explicit water and lipid molecules, but also reveal the transition pathways that are qualitatively consistent with previous CG and all-atom MD simulations.

### 2.2 A new case study: lipid transport through flippase ATP8A1-CDC50a

After validating the method, we used it to investigate the conformational transitions as well as the accompanying lipid transport process of the lipid flippase ATP8A1-CDC50a complex. ATP8A1 is one of the P4-ATPases, which functions with the regulatory protein CDC50a as the lipid flippase translocating the PS lipids from the exoplasmic leaflet of the plasma membrane to the cytoplasmic leaflet. ^44,45^ Like other P-type ATPases, P4-ATPases go through the Post-Albers cycle including E1, E1-ATP, E1P-ADP, E1P, E2P, and E2Pi-PL states during the lipid flipping process. ^46–48^ Among these states, the lipid translocation is mainly coupled to the transitions of E2Pi-PL-to-E1 states with the phosphate release at the P domain of P4-ATPases. Although the cryo-EM structures of these states have been obtained (Fig. 4a, b), the exquisite details of the conformational transitions between these functional states have not been revealed. ^46–48^ Thus, we utilized the Switching Gō-Martini approach to simulate the E2Pi-PL-to-E1 conformational transitions of the ATP8A1-CDC50a complex and investigated the lipid flipping process during the conformational changes.

**Figure 4:**
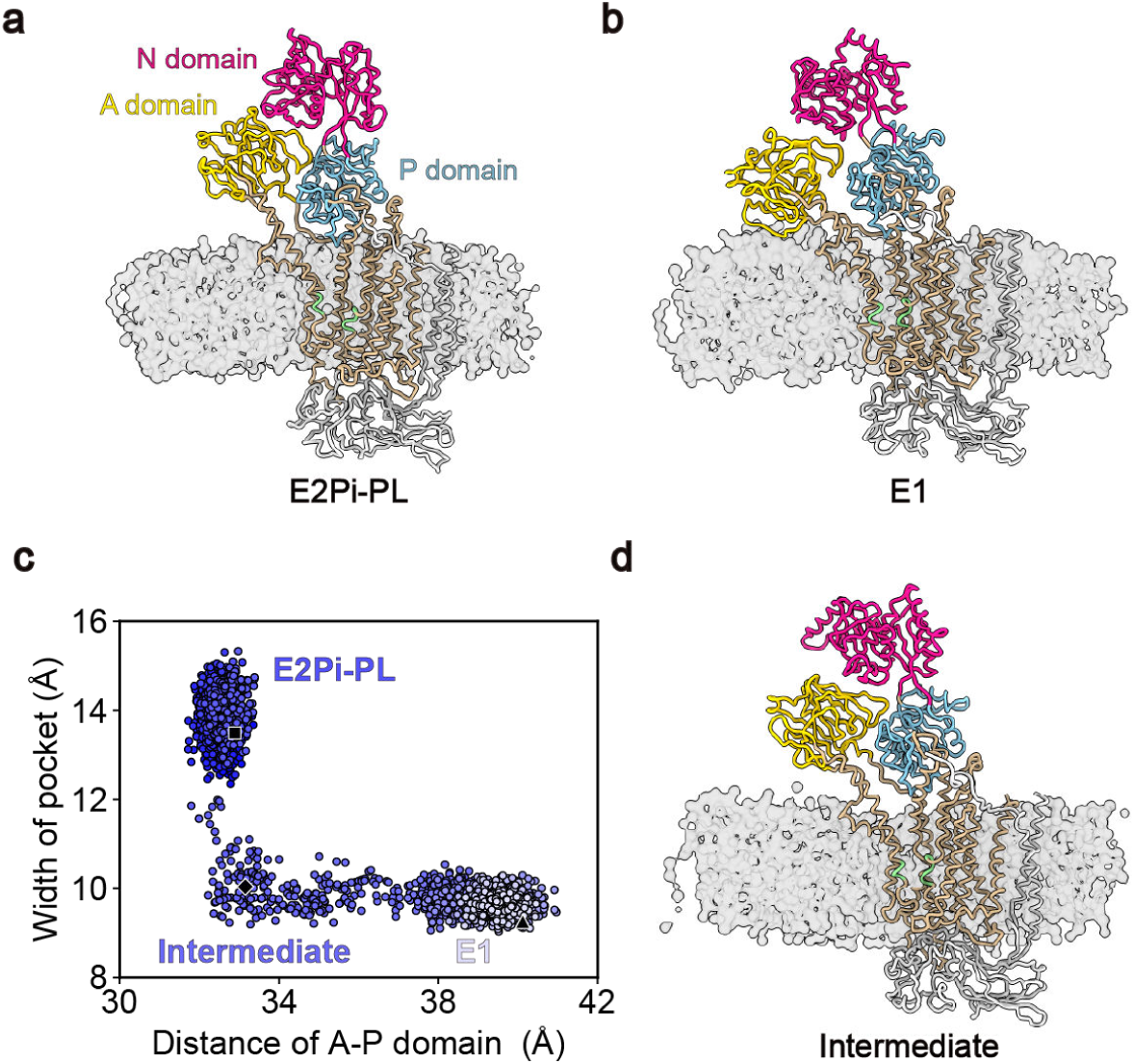
Switching Gō-Martini simulations for ATP8A1-CDC50a. **a, b, d,** Representative coarsegrained structures of ATP8A1-CDC50a in the E2Pi-PL, E1, and intermediate states embedded into the lipid bilayer, respectively. The protein is shown as the tan ribbon. The A, N, and P domains of ATP8A1 are labeled and colored gold, deep pink, and sky blue, respectively. The two helices constructing the window of the lipid-binding pocket are colored light green and CDC50a is colored light gray. The membrane is shown as gray surfaces. **c,** Two-dimension conformational transition pathway, as constructed with the distance of A-P domains and the width of the lipid-binding pocket. The blue color of the circles decreases along with the simulation time. The black square, diamond, and circle are representative of the E2Pi-PL, intermediate, and E1 states of ATP8A1-CDC50a.

#### 2.2.1 E2Pi-PL-to-E1 conformational transitions of ATP8A1-CDC50a

The CG model of the P4-ATPase complex in the E2Pi-PL state was constructed and embedded into an asymmetric membrane with POPS occupying 10% of the cytoplasmic leaflet and POPC occupying the remaining lipid bilayer (Fig. 4a). The PS lipid found in the binding pocket of ATP8A1 in the EM density map was also retained and transformed into the POPS CG model. A 200-ns simulation of ATP8A1-CDC50a in the E2Pi-PL state was performed followed by switching to the E1 state with a simulation of 800 ns. Totally 50 independent conformational transitions were obtained and RMSD of the complex during the main transition processes was shown in Fig. S4a. The RMSD with respect to the E1 state decreased from 7.6 ± 0.6 Å to 4.6 ± 0.7 Å after the switch operation, and meanwhile, the value with respect to the E2Pi-PL state increased from 3.6 ± 0.6 Å to 7.0 ± 0.6 Å. The changes of RMSD along with the switching simulation illustrated that the E2Pi-PL state of ATP8A1-CDC50a was successfully switched to the E1 state.

In a cell, the E2Pi-PL-to-E1 transition is facilitated by the phosphate release at the P domain of ATP8A1 with the A domain far away from it. In this process, the lipid-binding pocket of ATP8A1 closes and the bound PS lipid can be translocated from the exoplasmic leaflet to the cytoplasmic leaflet of the plasma membrane. Thus, the distance between the A domain and the P domain of ATP8A1 and the width of the binding pocket were used as reaction coordinates to monitor the E2Pi-PL-to-E1 conformational transitions of ATP8A1-CDC50a. As shown in Fig. 4c, d and Fig. S5, the width of the binding pocket, measured as the distance between two helices (residues 100-104 and 878-883), first decreased from 14.0 ± 0.4 Å to 9.8 ± 0.2 Å. Then, the distance of A–P domains had a significant increase from 32.6 ± 0.3 Å to 39.6 ± 0.4 Å. Upon analyzing all 50 simulations, we found that 36 repeats (72%) out of our simulations followed the aforementioned pathway, while a distinct conformational transition was observed in 5 repeats (10%) in which the lipid-binding pocket underwent closure and the A domain simultaneously moved away from the P domain (Fig. S4a–d). The rest nine repeats (18%) appeared to follow the first pathway but were incomplete and remained in the intermediate state in our simulation time (Fig. S4e, f), possibly due to the limited simulation time.

#### 2.2.2 Lipid translocation mechanism of ATP8A1-CDC50a

Along with the conformational transition, a detailed analysis of the lipid translocation process was conducted. In 23 out of 50 independent simulations, complete lipid translocation events were observed (Fig. 5a), wherein the bound POPS lipid was found to be flipped from the exoplasmic leaflet to the cytoplasmic leaflet. In 13 out of the 50 simulations, the bound lipid failed to complete the flipping process and returned to the exoplasmic leaflet of the membrane (Fig. S6a). The remaining 14 simulations were incomplete, in which the lipid was trapped in the binding pocket of ATP8A1 (Fig. S6b). Extending the simulations for an additional 1000 ns resulted in more flipping events being observed (6 repeats completed and 2 failed), indicating that the lipid trapping state is metastable and a majority of the bound POPS tend to flip during the E2Pi-PL-to-E1 transition (Fig. S6b).

**Figure 5:**
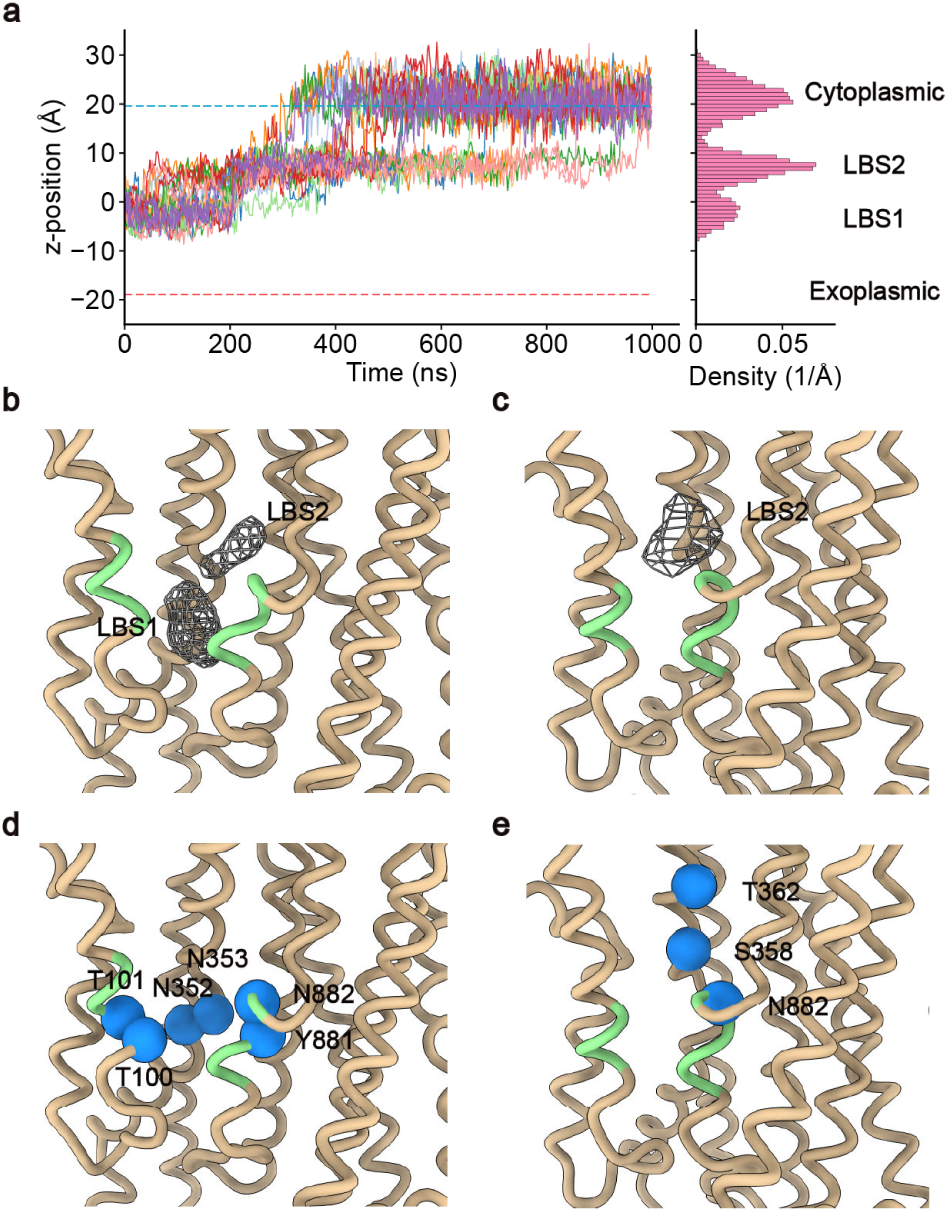
Lipid translocation and lipid-binding sites of ATP8A1-CDC50a. **a,** Traces (left) and distribution density (right) of the lipid heads across the z-axis in the course of simulations. All lipids successfully translocated in our 50 simulations are shown as solid lines with different colors. Red and blue dash lines represent the exoplasmic and cytoplasmic leaflets of the bilayer membrane. The positions of lipid binding sites (LBS1, LBS2) are also labeled. **b, c,** Lipid binding sites for E2Pi-PL and E1 states of ATP8A1-CDC50a, respectively. The protein complex is shown as the tan ribbon, with the helices forming the window of the lipid-binding pocket colored light green. Lipid binding sites are shown as the mesh representation. **d, e,** The polar residues around Lipid binding site 1 and 2 for E2Pi-PL and E1 states of ATP8A1-CDC50a, respectively. The BB atoms of the polar residues are shown as dodger blue spheres.

The lipid head density maps of the bound POPS were used to reveal the lipid binding site during the translocation process. In the E2Pi-PL and E1 states, two binding sites were identified as shown in Fig. 5b, c. Lipid binding site 1 was located in the same binding pocket as found by the cryo-EM density maps, which was surrounded by residues T100, T101, P104, N352, N353, I357, I878, G879, L880, Y881, and N882 (Fig. 5d). Lipid binding site 2 was formed by residues F107, V111, S358, V361, T362, N882, V883, A887, and L891, which was identified in our simulations (Fig. 5e). The density maps of the two lipid binding sites indicated that the bound lipid could move between the two binding sites in the E2Pi-PL state, while mainly bound to Lipid binding site 1 (Fig. 5b). When we switched off the E2Pi-PL state and switched on the E1 topology, the lipid in Lipid binding site 1 was squeezed and transported to Lipid binding site 2 (movie S1). Then, the tail of the bound lipid changed its direction from perpendicular to parallel to the membrane surface (Fig. 6a, b). With the simulation going, a majority of the bound lipids flipped to the cytoplasmic leaflet of the membrane (Fig. 6c), although some of them returned to the exoplasmic leaflet of the membrane.

**Figure 6:**
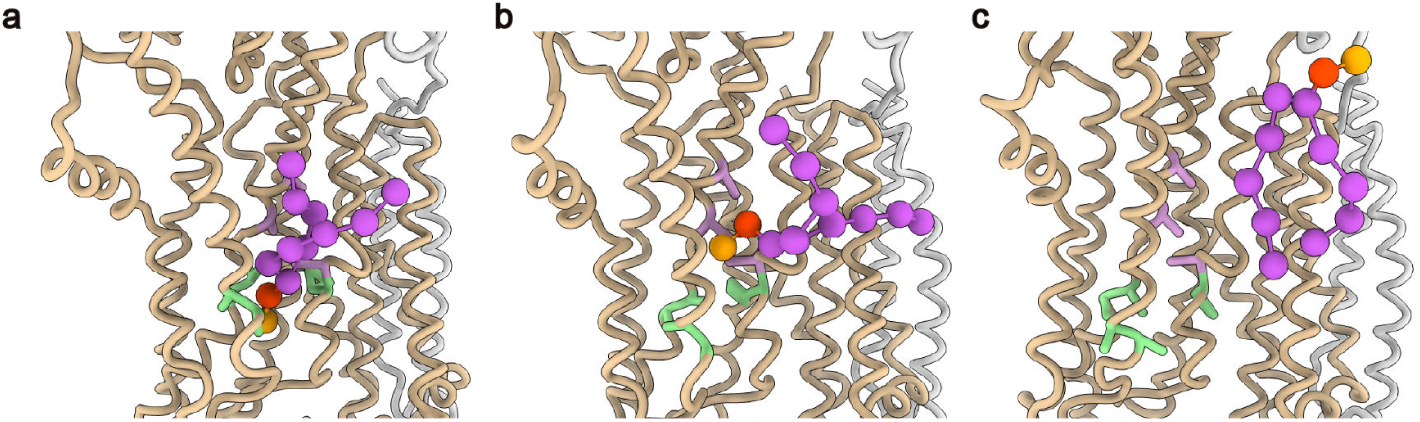
Lipid translocation process of ATP8A1-CDC50a. **a, b, c,** Zoom-in views of lipid translocation process of ATP8A1-CDC50a with the bound lipid in Lipid-binding site 1 (**a**), Lipid-binding site 2 (**b**), and cytoplasmic leaflet of the plasma membrane (**c**), respectively. The protein is shown as a ribbon with ATP8A1 colored tan and CDC50a colored light gray. The POPS lipid is shown as the ball-and-stick model with the tail colored medium orchid and the lipid head group colored orange and red. The essential polar residues in Lipid-binding site 1 and 2 are colored light green and plum, respectively, with the side chains shown as a stick.

As shown in Fig. 5d, Lipid binding site 1 is surrounded by the polar residues including T100, T101, N352, N353, Y881, and N882, which can selectively bind to certain types of lipids (PS for ATP8A1), ^46^ while Lipid binding site 2 plays a role as the intermediate harbor that is important for the translocation of the bound lipid. Interestingly, the density blobs at the same location of Lipid binding site 2 were reported in the cryo-EM structures from different types of P4-ATPases. ^49^ Additionally, Lipid binding site 2 is composed of some polar residues including S358, T362, and N882 (Fig. 5e). Mutations of these three residues in the homology proteins were reported to significantly decrease the transportation ability. The S365A and T369A mutations of ATP8A2 (S358 and T362 of ATP8A1) displayed significant reductions in the ATPase activity. ^50^ The N1226A mutation of Dnf1 (corresponding to N882 of ATP8A1) could completely halt the transportation of lipids even though the mutant protein was still localized on the plasma membrane. ^47^ And more importantly, the S403Y mutation in ATP8B1 (equivalent to S358 of ATP8A1) was found in the patients with progressive familial intrahepatic cholestasis type 1 (PFIC1). ^51^ These experimental results support that Lipid binding site 2 plays an essential role in the lipid translocation process. All these findings demonstrate that our Switching Gō-Martini simulations effectively captured the E2Pi-PL-to-E1 conformational transition of ATP8A1-CDC50a and shed new light on the lipid translocation mechanism of the P4-ATPase flippase.

## 3 Discussion

Traditional Gō-like CG models have obtained great success in the simulation of protein conformational transition. However, the development of these models has been impeded by several limitations: (1) the implicit solvent, which cannot properly reveal the interaction between proteins and water or charged ions; (2) the implicit membrane environment, which cannot show the association and disassociation of lipids around proteins. Martini model utilizes explicit solvent and develops a great diversity of lipids, providing an important computational tool for the understanding of interactions between proteins and lipids, and has therefore been extensively utilized in this kind of studies. ^18,19,52,53^ However, the elastic network restraints in the previous martini protein model used to stabilize the second or higher structures impede the conformational changes of proteins. Recently, the Gō-like martini model was developed with Lennard-Jones potential replacing the harmonic potential in the elastic network, which showed similar dynamics of proteins compared with the all-atom model. ^21^ And the Gō-like martini model can simulate the folding process of small proteins and the unfolding process induced by the external pulling force. ^21^ The release of the latest Martini 3 further facilitate the development in this direction, as demonstrated in the study of a small molecular switch recently. ^23^ These encouraging studies inspired us to implement the Gō-like Martini model to capture the conformational changes of multi-state proteins.

Therefore, here we develop the Switching Gō-Martini method and test its ability to sample the conformational transitions of proteins. Three well-tested systems, from simple to frustrated and from soluble to membrane-embedded, have validated that the method can correctly capture the conformational changes of proteins without fine-tuning parameters. Moreover, utilizing the explicit-membrane models in the Martini force field, we were able to simulate the lipid-protein interaction in the lipid translocation process of the P4-ATPases. A series of 50 independent simulations employing the Switching Gō-Martini method revealed the exquisite details of the conformational transitions of ATP8A1-CDC50a from the E2Pi-PL state to the E1 state. The entire process of lipid translocation during these conformational changes was captured for the first time and the underlying mechanism of P4-ATPase-mediated lipid translocation was elucidated (Fig. 7). This is a significant advantage of the Switching Gō-Martini method in comparison to other existing Gō-like methods.

**Figure 7:**
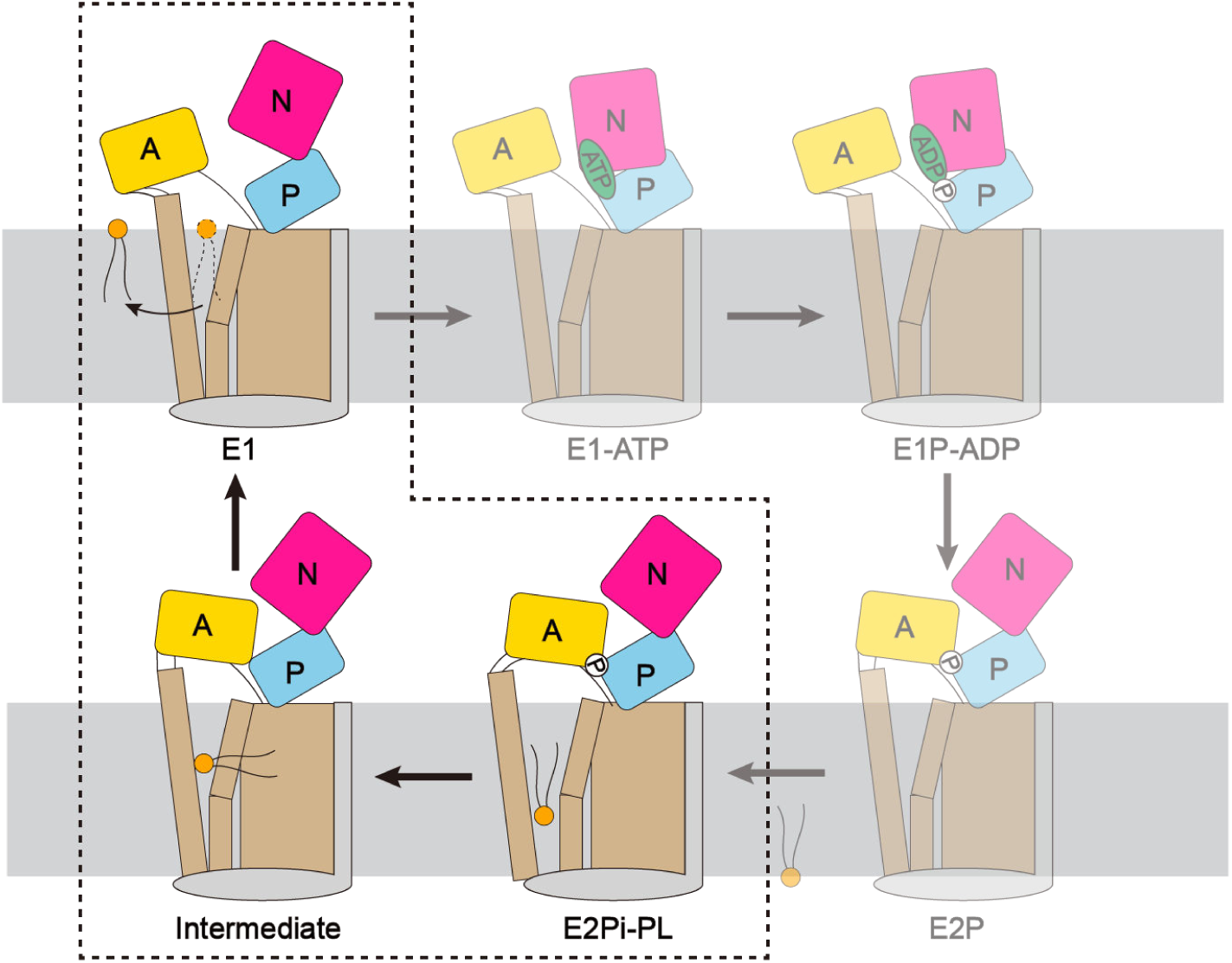
Lipid translocation mechanism of the P4-ATPase flippases. Schematic model of the lipid translocation cycle for the P4-ATPase flippases. The model is depicted with the same color codes as in Fig. 4. The E2Pi-PL-to-E1 conformational transition conducted here is enclosed by the black dashed box. The other conformational transitions that were not simulated are depicted as transparent. At the beginning of the lipid translocation process, Lipid binding site 1 of the P4-ATPase flippase in the E2Pi-PL state specifically attracts and binds a phospholipid. Subsequently, with the phosphate release at the P domain of the P4-ATPase, Lipid binding site 1 undergoes closure, resulting in the displacement of the bound lipid towards Lipid binding site 2. Then the phospholipid leaves Lipid binding site 2 and translocates towards the inner leaflet of the membrane with the assistance of thermal fluctuations, and the flippase adopts the E1 conformation, ready to initiate another phosphorylation and dephosphorylation cycle.

Although the approach has been successful in simulating conformational transitions of proteins, the method cannot accurately depict the free energy landscape between two or more states, due to the so-called “vertical excitation” of proteins when switching the state. This intrinsic problem pertains to the Gō-like models employing the switching approach. ^17^ In some cases, proteins might fall into the final state through an inappropriate pathway due to the overestimated energy for overcoming the transition states (Fig. 1a). For example, in the AdK simulations, 21.4% of our simulations utilized the middle pathway in which NMP and LID domains opened concurrently (Fig. S2f). This specific phenomenon had not been previously reported in other CG simulations, ^15,16^ where only the NMP-closing pathway and LID-closing pathway were observed. The discrepancy could potentially be attributed to the switching approach in our simulations, allowing proteins with higher energy levels to adopt improper pathways. Therefore, a multiple-basin Gō-Martini method would be desired in future studies, which can be used to obtain more spontaneous and accurate transitions between different states of proteins. ^14,15,54,55^

## 4 Materials and Methods

### 4.1 Simulation settings

All simulations in our work were performed using the coarse-grained (CG) force field Martini 3^19^ (version 3.0.0) with the MD package Gromacs^56^ (version 2020.x or 2021.x). Details about generating protein CG models and setting up simulation systems are given in the following sections. For the MD simulations, the systems were minimized and then equilibrated in the NPT ensemble with position restraints (1,000 *kJ/mol/nm*^2^) on the backbones of CG proteins. Productions were conducted at a constant isotropic or semi-isotropic pressure of 1 atm and a temperature of 310.15 K according to the systems. The V-rescale algorithm ^57^ with a coupling time of 1.0 ps was used to maintain the temperature of the systems and the Parrinello-Rahman thermostat ^58^ with a coupling time of 12.0 ps was performed to maintain the pressure of the systems. The electrostatic interaction and the van der Waals interaction were both cut off at 1.1 nm and the long-range electrostatic interaction was computed by using the reaction-field method ^59^ with a relative dielectric constant (epsilon_r = 15).

The switching method for conformational transitions of different states of proteins is introduced here. Briefly, the systems were first simulated to oscillate around State A by using the topology of State A. Before completely switching to State B, the systems must be relaxed to fit the new topology by performing a short and slow simulation (1,000 steps × 0.002 ps per step) using the topology of State B. Then production simulations were conducted using the topology of State B, during which the systems could be gradually transited to State B. All simulation parameters for running CG systems are listed in Supplementary Table S1, and the results of transition pathways are shown in Supplementary Table S2. In the Supplementary Materials, we provide a tutorial to show how to set up such simulations, using the active-to-inactive transition of β2AR as an example.

### 4.2 Glutamine binding protein CG model

The open- and closed-state CG models of glutamine binding protein (GlnBP) were built using the Martini 3 force field and the structures with the PDB codes 1GGG ^28^ and 1WDN, ^27^ respectively. The sequences of both proteins were trimmed to have the same length (residues 5-224). The topologies of the open- and closed-states of GlnBP were generated using the program martinize2.py (https://github.com/marrink-lab/vermouth-martinize). Additional Lennard-Jones interactions for the Gō-like model were generated based on the OV and rCSU contacts (http://info.ifpan.edu.pl/~rcsu/rcsu/index.html) and set up using the program create_goVirt.py (http://md.chem.rug.nl) with a distance cutoff of 0.3-1.1 nm and a dissociation energy of *ε* = 12.0 kJ/mol. The open state of GlnBP was chosen as the initial state in our simulations. The CG protein and water were placed in a rectangular box (8×8×7 *nm*^3^) using the program insane.py. ^60^ 150 mM NaCl was added to keep the whole system charge neutral. 50 CG simulations with 300 × 3 ns were performed to investigate the open-closed-open conformational transition path of GlnBP. Additionally, to further test the robustness of the method, one simulation comprising 10 cycles of open↔closed transitions was also performed with the same parameters.

### 4.3 Adenylate kinase CG model

The open- and closed-state CG models of adenylate kinase (AdK) were built using the Martini 3 force field and the structures with the PDB codes 4AKE^33^ and 1ANK, ^32^ respectively. The topologies of the open- and closed-states of AdK were generated using the program martinize2.py (https://github.com/marrink-lab/vermouth-martinize). Additional Lennard-Jones interactions for the Gō-like model were generated based on the OV and rCSU contacts (http://info.ifpan.edu.p l/~rcsu/rcsu/index.html) and set up using the program create_goVirt.py (http://md.chem.rug.nl) with a distance cutoff of 0.3-1.1 nm and a dissociation energy of *ε* = 12.0 kJ/mol. The closed state of AdK was chosen as the initial state in our simulations. The CG protein and water were placed in a rectangular box (7 × 9 × 9 *nm*^3^) using the program insane.py. ^60^ 150 mM NaCl was added to keep the whole system charge neutral. 70 CG simulations with 100 × 2 ns were performed to investigate the closed-to-open conformational transition path of AdK.

### 4.4 *β*_2_-adrenergic receptor CG model

The active- and inactive-state CG models of *β*_2_-adrenergic receptor (β2AR) were built using the Martini 3 force field and the structures with the PDB codes 3SN6^40^ and 2RH1, ^41^ respectively. The missing loop and mutations (residues 176-178, mutations M96T and M98T) of 3SN6 were repaired using Modeller. ^61^ The sequences of both proteins were trimmed to have the same length (residues 30-230 and 265-341). The topologies of the active and inactive states of β2AR were generated using the program martinize2.py (https://github.com/marrink-lab/vermouth-mar-tinize). Additional Lennard-Jones interactions for the Gō-like model were generated based on the OV and rCSU contacts (http://info.ifpan.edu.pl/~rcsu/rcsu/index.html) and set up using the program create_goVirt.py (http://md.chem.rug.nl) with a distance cutoff of 0.3-1.1 nm and a dissociation energy of *ε* = 12.0 kJ/mol. The active state of β2AR was chosen as the initial state in our simulations. The CG protein was embedded into a POPC bilayer within a rectangular box (8 × 8 × 10 *nm*^3^) using the program insane.py. ^60^ 150 mM NaCl was added to keep the whole system charge neutral. 40 CG simulations with 500 × 2 ns were performed to investigate the active-to-inactive conformational transition path of β2AR.

### 4.5 The P4-ATPase flippase ATP8A1-CDC50a CG model

The CG models of the E2Pi-PL and E1 states of the P4 ATPase ATP8A1-CDC50a complex were built using the Martini 3 force field and the structures with the PDB codes 6K7M and 6K7G, respectively. ^46^ The missing loops (residues 237-242 and 434-445) of ATP8A1 were repaired using Modeller. ^61^ The sequences of both structures were trimmed to have the same length (residues 35-696 and 725-1063 of ATP8A1 and residues 27-350 of CDC50a). The topologies of the E2Pi-PL and E1 states of ATP8A1-CDC50a were generated using the program martinize2.py (https://github.com/marrink-lab/vermouth-martinize). Additional Lennard-Jones interactions for the Gō-like model were generated based on the OV and rCSU contacts (http://info.ifpan.edu.pl/~rcsu/rcsu/index.html), which were set up using the program create_goVirt_for_multimer.py modified based on the initial program create_goVirt.py (http://md.chem.rug.nl) by ourselves, with a distance cutoff of 0.3-1.1 nm and a dissociation energy of *ε* = 12.0 kJ/mol. The E2Pi-PL state of ATP8A1-CDC50a was chosen as the initial state in our simulations. And the binding PS lipid found in the cryo-EM structure was also retained and converted to the POPS CG model, while the phosphate ligand was considered implicit and not displayed in our simulations. The CG protein and the bound lipid were embedded into an asymmetric membrane within a rectangular box (12 × 12 × 19 *nm*^3^) using the program insane.py, ^60^ in which POPS occupied 10% of the cytoplasmic leaflet with POPC occupying the remaining membrane. 150 mM NaCl was added to keep the whole system charge neutral. 50 CG simulations of 200 ns (E2Pi-PL) plus 800 ns (E1) were performed to investigate the E2Pi-PL→E1 conformational transition path and the lipid translocation process of the P4-ATPase ATP8A1-CDC50a complex. For those in which the lipid was trapped in the binding pocket of ATP8A1, an additional 1000 ns was conducted to investigate the stability of the lipid trapping state.

### 4.6 Classification of AdK conformational transition pathways

Conformational transition paths of AdK were classified as LID-closing pathway, NMP-closing pathway, and middle pathway by using the combination of two angles *θ_NMP_* and *θ_LID_*. The angles were defined according to the center of mass of the following atoms: BB atoms of residues 35-55, 90-100, 115-125 for *θ_NMP_*, and BB atoms of residues 125-153, 115-125, 204-212 for *θ_LID_*. AdK transitions were classified as LID-closing pathways if *θ_NMP_* of transition states was more than 70 degrees and *θ_LID_* was less than 90 degrees, NMP-closing pathway if *θ_NMP_* of transition states was less than 50 degrees and *θ_LID_* was more than 105 degrees, or middle pathway for those not belong to LID/NMP-closing pathway. The representative intermediate structures were obtained by using the k-means clustering method.

### 4.7 Classification of β2AR conformational transition pathways

Conformational transition paths of β2AR deactivation were classified as normal pathway and abnormal pathways by using the combination of helix 6–helix 3 distance and the RMSD of NPxxY motif (residues 322-327) aligned to the inactive state. Helix 6–helix 3 distance was measured as the distance between R131 and L272 BB atoms. β2AR transitions were classified as normal pathway if the helix 6–helix 3 distance of transition states was more than 12 Å and the RMSD of NPxxY motif was less than 0.6 Å, or abnormal pathway if the helix 6–helix 3 distance of transition states was less than 12 Å and the RMSD of NPxxY motif was more than 0.6 Å. The representative intermediate structures were obtained by using the k-means clustering method.

### 4.8 Classification of ATP8A1-CDC50a conformational transition pathways

Conformational transition paths of ATP8A1-CDC50a were classified as main pathway and middle pathway by using the combination of the distance of A-P domains and the width of the lipid-binding pocket. The distance of A-P domains was measured as the distance between the center of mass of BB atoms of the A domain (residues 35-51 and 134-279) and P domain (residues 384-416 and 649-819). The width of the lipid-binding pocket was calculated as the distance between the two helices (residues 100-104 and 878-883) constructing the window of the lipid-binding pocket. ATP8A1-CDC50a transitions were classified as main pathway if the distance of A-P domains of transition states was less than 34 Å and the width of the pocket was less than 11 Å, or middle pathway if the disassociation of A-P domains and the closure of the lipid-binding pocket happened simultaneously. The representative intermediate structures were obtained by using the k-means clustering method.

### 4.9 Simulation Analysis

The lipid head density was obtained by computing the occupancy of the lipid heads of the bound POPS in the three-dimensional space using the Volmap plugin of VMD. ^62^ The grid points had a spacing distance of 0.1 nm. The polar residues around the binding sites were obtained by analyzing the interaction between the bound lipid heads and residues with a distance cutoff of 6 Å and an occupancy cutoff of 40%. Additionally, geometry calculation, RMSD, and protein-lipid interaction were analyzed with MDAnalysis. ^63^ All structural figures and the movie were generated using ChimeraX. ^64^

## Acknowledgements

We thank Dr. Yuji Sugita and Dr. Chigusa Kobayashi at RIKEN for insightful suggestions provided through numerous discussions, and we thank Dr. Long Li at Peking University for critical reading of the manuscript. This work was supported by the National Key R&D Program of China (2021YFE0108100).

## Author Contributions

S.Y. and C.S. conceived the idea and designed the research. S.Y. conducted simulations and analyzed data. S.Y. and C.S. wrote the manuscript. C.S. acquired funding and supervised the work.

## References

[1] Tam, R.; Saier, M. H. Structural, functional, and evolutionary relationships among extracellular solute-binding receptors of bacteria. Microbiological Reviews 1993, 57, 320–346.

[2] Dzeja, P.; Terzic, A. Adenylate kinase and AMP signaling networks: metabolic monitoring, signal communication and body energy sensing. International Journal of Molecular Sciences 2009, 10, 1729–1772.

[3] Kefauver, J. M.; Ward, A. B.; Patapoutian, A. Discoveries in structure and physiology of mechanically activated ion channels. Nature 2020, 587, 567–576.

[4] Philipson, L. H. β-agonists and metabolism. Journal of Allergy and Clinical Immunology 2002, 110, S313–S317.

[5] Wickstrand, C.; Nogly, P.; Nango, E.; Iwata, S.; Standfuss, J.; Neutze, R. Bacteriorhodopsin: Structural insights revealed using x-ray lasers and synchrotron radiation. Annual Review of Biochemistry 2019, 88, 59–83.

[6] Stiller, J. B.; Otten, R.; Häussinger, D.; Rieder, P. S.; Theobald, D. L.; Kern, D. Structure determination of high-energy states in a dynamic protein ensemble. Nature 2022, 603, 528–535.

[7] Zhang, S.; Zou, S.; Yin, D.; Zhao, L.; Finley, D.; Wu, Z.; Mao, Y. USP14-regulated allostery of the human proteasome by time-resolved cryo-EM. Nature 2022, 605, 567–574.

[8] Hoffer, N. Q.; Woodside, M. T. Probing microscopic conformational dynamics in folding reactions by measuring transition paths. Current Opinion in Chemical Biology 2019, 53, 68–74.

[9] Sugita, Y.; Okamoto, Y. Replica-exchange molecular dynamics method for protein folding. Chemical Physics Letters 1999, 314, 141–151.

[10] Laio, A.; Parrinello, M. Escaping free-energy minima. Proceedings of the National Academy of Sciences 2002, 99, 12562–12566.

[11] Torrie, G. M.; Valleau, J. P. Nonphysical sampling distributions in Monte Carlo free-energy estimation: Umbrella sampling. Journal of Computational Physics 1977, 23, 187–199.

[12] Husic, B. E.; Pande, V. S. Markov State Models: From an Art to a Science. Journal of the American Chemical Society 2018, 140, 2386–2396.

[13] Takada, S. Gō model revisited. Biophysics and physicobiology 2019, 16, 248–255.

[14] Okazaki, K. I.; Koga, N.; Takada, S.; Onuchic, J. N.; Wolynes, P. G. Multiple-basin energy landscapes for large-amplitude conformational motions of proteins: Structure-based molecular dynamics simulations. Proceedings of the National Academy of Sciences of the United States of America 2006, 103, 11844–11849.

[15] Lu, Q.; Wang, J. Single molecule conformational dynamics of adenylate kinase: Energy landscape, structural correlations, and transition state ensembles. Journal of the American Chemical Society 2008, 130, 4772–4783.

[16] Shinobu, A.; Kobayashi, C.; Matsunaga, Y.; Sugita, Y. Coarse-Grained Modeling of Multiple Pathways in Conformational Transitions of Multi-Domain Proteins. Journal of Chemical Information and Modeling 2021, 61, 2427–2443.

[17] Koga, N.; Takada, S. Folding-based molecular simulations reveal mechanisms of the rotary motor F1-ATPase. Proceedings of the National Academy of Sciences of the United States of America 2006, 103, 5367–5372.

[18] Marrink, S. J.; Risselada, H. J.; Yefimov, S.; Tieleman, D. P.; Vries, A. H. D. The MARTINI force field: Coarse grained model for biomolecular simulations. Journal of Physical Chemistry B 2007, 111, 7812–7824.

[19] Souza, P. C. et al. Martini 3: a general purpose force field for coarse-grained molecular dynamics. Nature Methods 2021, 18, 382–388.

[20] Negami, T.; Shimizu, K.; Terada, T. Coarse-grained molecular dynamics simulation of protein conformational change coupled to ligand binding. Chemical Physics Letters 2020, 742, 137144.

[21] Poma, A. B.; Cieplak, M.; Theodorakis, P. E. Combining the MARTINI and Structure-Based Coarse-Grained Approaches for the Molecular Dynamics Studies of Conformational Transitions in Proteins. Journal of Chemical Theory and Computation 2017, 13, 1366–1374.

[22] Mahmood, M. I.; Poma, A. B.; Okazak, K. I. Optimizing Gō-MARTINI coarse-grained model for F-BAR protein on Lipid membrane. Frontiers in Molecular Biosciences 2021, 8, 619381.

[23] Vainikka, P.; Marrink, S. J. Martini 3 Coarse-Grained Model for Second-Generation Unidirectional Molecular Motors and Switches. Journal of Chemical Theory and Computation 2023, 19, 596–604, PMID: 36625495.

[24] Hyeon, C.; Lorimer, G. H.; Thirumalai, D. Dynamics of allosteric transitions in GroEL. Proceedings of the National Academy of Sciences of the United States of America 2006, 103, 18939–18944.

[25] Yao, X. Q.; Kenzaki, H.; Murakami, S.; Takada, S. Drug export and allosteric coupling in a multidrug transporter revealed by molecular simulations. Nature Communications 2010, 1, 1–8.

[26] Brandani, G. B.; Takada, S. Chromatin remodelers couple inchworm motion with twist-defect formation to slide nucleosomal DNA. PLoS Computational Biology 2018, 14, e1006512.

[27] Sun, Y. J.; Rose, J.; Wang, B. C.; Hsiao, C. D. The structure of glutamine-binding protein complexed with glutamine at 1.94 Å resolution: Comparisons with other amino acid binding proteins. Journal of Molecular Biology 1998, 278, 219–229.

[28] Hsiao, C. D.; Sun, Y. J.; Rose, J.; Wang, B. C. The crystal structure of glutamine-binding protein from Escherichia coli. Journal of Molecular Biology 1996, 262, 225–242.

[29] Zhang, L.; Wu, S.; Feng, Y.; Wang, D.; Jia, X.; Liu, Z.; Liu, J.; Wang, W. Ligand-bound glutamine binding protein assumes multiple metastable binding sites with different binding affinities. Communications Biology 2020, 3, 419.

[30] Kooshapur, H.; Ma, J.; Tjandra, N.; Bermejo, G. A. NMR Analysis of Apo Glutamine-Binding Protein Exposes Challenges in the Study of Interdomain Dynamics. Angewandte Chemie - International Edition 2019, 58, 16899–16902.

[31] Feng, Y.; Zhang, L.; Wu, S.; Liu, Z.; Gao, X.; Zhang, X.; Liu, M.; Liu, J.; Huang, X.; Wang, W. Conformational Dynamics of apo-GlnBP Revealed by Experimental and Computational Analysis. Angewandte Chemie - International Edition 2016, 55, 13990–13994.

[32] Berry, M. B.; Meador, B.; Bilderback, T.; Liang, P.; Glaser, M.; Phillips, G. N. The closed conformation of a highly flexible protein: The structure of E. coli adenylate kinase with bound AMP and AMPPNP. Proteins: Structure, Function, and Bioinformatics 1994, 19, 183–198.

[33] Müller, C. W.; Schlauderer, G. J.; Reinstein, J.; Schulz, G. E. Adenylate kinase motions during catalysis: An energetic counterweight balancing substrate binding. Structure 1996, 4, 147–156.

[34] Henzler-Wildman, K. A.; Thai, V.; Lei, M.; Ott, M.; Wolf-Watz, M.; Fenn, T.; Pozharski, E.; Wilson, M. A.; Petsko, G. A.; Karplus, M.; Hübner, C. G.; Kern, D. Intrinsic motions along an enzymatic reaction trajectory. Nature 2007, 450, 838–844.

[35] Li, W.; Wolynes, P. G.; Takada, S. Frustration, specific sequence dependence, and nonlinearity in large-amplitude fluctuations of allosteric proteins. Proceedings of the National Academy of Sciences of the United States of America 2011, 108, 3504–3509.

[36] Stiller, J. B.; Kerns, S. J.; Hoemberger, M.; Cho, Y. J.; Otten, R.; Hagan, M. F.; Kern, D. Probing the transition state in enzyme catalysis by high-pressure NMR dynamics. Nature Catalysis 2019, 2, 726–734.

[37] Seyler, S. L.; Beckstein, O. Sampling large conformational transitions: Adenylate kinase as a testing ground. Molecular Simulation 2014, 40, 855–877.

[38] Kobayashi, C.; Matsunaga, Y.; Koike, R.; Ota, M.; Sugita, Y. Domain Motion Enhanced (DoME) Model for Efficient Conformational Sampling of Multidomain Proteins. Journal of Physical Chemistry B 2015, 119, 14584–14593.

[39] Barisione, G.; Baroffio, M.; Crimi, E.; Brusasco, V. Beta-adrenergic agonists. Pharmaceuticals 2010, 3, 1016–1044.

[40] Rasmussen, S. G. et al. Crystal structure of the *β* 2 adrenergic receptor-Gs protein complex. Nature 2011, 477, 549–557.

[41] Cherezov, V.; Rosenbaum, D. M.; Hanson, M. A.; Rasmussen, S. G.; Foon, S. T.; Kobilka, T. S.; Choi, H. J.; Kuhn, P.; Weis, W. I.; Kobilka, B. K.; Stevens, R. C. High-resolution crystal structure of an engineered human *β* 2-adrenergic G protein-coupled receptor. Science 2007, 318, 1258–1265.

[42] Dror, R. O.; Arlow, D. H.; Maragakis, P.; Mildorf, T. J.; Pan, A. C.; Xu, H.; Borhani, D. W.; Shaw, D. E. Activation mechanism of the *β* 2-adrenergic receptor. Proceedings of the National Academy of Sciences of the United States of America 2011, 108, 18684–18689.

[43] Kohlhoff, K. J.; Shukla, D.; Lawrenz, M.; Bowman, G. R.; Konerding, D. E.; Belov, D.; Altman, R. B.; Pande, V. S. Cloud-based simulations on Google Exacycle reveal ligand modulation of GPCR activation pathways. Nature Chemistry 2014, 6, 15–21.

[44] Montigny, C.; Lyons, J.; Champeil, P.; Nissen, P.; Lenoir, G. On the molecular mechanism of flippase- and scramblase-mediated phospholipid transport. Biochimica et Biophysica Acta-Molecular and Cell Biology of Lipids 2016, 1861, 767–783.

[45] López-Marqués, R. L.; Gourdon, P.; Pomorski, T. G.; Palmgren, M. The transport mechanism of P4 ATPase lipid flippases. Biochemical Journal 2020, 477, 3769–3790.

[46] Hiraizumi, M.; Yamashita, K.; Nishizawa, T.; Nureki, O. Cryo-EM structures capture the transport cycle of the P4-ATPase flippase. Science 2019, 365, 1149–1155.

[47] Bai, L.; You, Q.; Jain, B. K.; Duan, H. D.; Kovach, A.; Graham, T. R.; Li, H. Transport mechanism of P4 ATPase phosphatidylcholine flippases. eLife 2020, 9, e62163.

[48] Xu, J.; He, Y.; Wu, X.; Li, L. Conformational changes of a phosphatidylcholine flippase in lipid membranes. Cell Reports 2022, 38, 110518.

[49] He, Y.; Xu, J.; Wu, X.; Li, L. Structures of a P4-ATPase lipid flippase in lipid bilayers. Protein and Cell 2020, 11, 458–463.

[50] Vestergaard, A. L.; Coleman, J. A.; Lemmin, T.; Mikkelsen, S. A.; Molday, L. L.; Vilsen, B.; Molday, R. S.; Peraro, M. D.; Andersen, J. P. Critical roles of isoleucine-364 and adjacent residues in a hydrophobic gate control of phospholipid transport by the mammalian P4-ATPase ATP8A2. Proceedings of the National Academy of Sciences of the United States of America 2014, 111, E1334–1343.

[51] Klomp, L. W. et al. Characterization of mutations in ATP8B1 associated with hereditary cholestasis. Hepatology 2004, 40, 27–38.

[52] Buyan, A.; Cox, C. D.; Barnoud, J.; Li, J.; Chan, H. S.; Martinac, B.; Marrink, S. J.; Corry, B. Piezo1 Forms Specific, Functionally Important Interactions with Phosphoinositides and Cholesterol. Biophysical Journal 2020, 119, 1683–1697.

[53] Mosalaganti, S.; Obarska-Kosinska, A.; Siggel, M.; Taniguchi, R.; Turoňová, B.; Zimmerli, C. E.; Buczak, K.; Schmidt, F. H.; Margiotta, E.; Mackmull, M. T.; Hagen, W. J.; Hummer, G.; Kosinski, J.; Beck, M. AI-based structure prediction empowers integrative structural analysis of human nuclear pores. Science 2022, 376, eabm9506.

[54] Best, R. B.; Chen, Y. G.; Hummer, G. Slow protein conformational dynamics from multiple experimental structures: The helix/sheet transition of Arc repressor. Structure 2005, 13, 1755–1763.

[55] Shinobu, A.; Kobayashi, C.; Matsunaga, Y.; Sugita, Y. Coarse-Grained Modeling of Multiple Pathways in Conformational Transitions of Multi-Domain Proteins. Journal of Chemical Information and Modeling 2021, 61, 2427–2443.

[56] Abraham, M. J.; Murtola, T.; Schulz, R.; Páll, S.; Smith, J. C.; Hess, B.; Lindah, E. Gromacs: High performance molecular simulations through multi-level parallelism from laptops to supercomputers. SoftwareX 2015, 1-2, 19–25.

[57] Bussi, G.; Donadio, D.; Parrinello, M. Canonical sampling through velocity rescaling. Journal of Chemical Physics 2007, 126, 014101.

[58] Parrinello, M.; Rahman, A. Polymorphic transitions in single crystals: A new molecular dynamics method. Journal of Applied Physics 1981, 52, 7182–7190.

[59] Tironi, I. G.; Sperb, R.; Smith, P. E.; Gunsteren, W. F. V. A generalized reaction field method for molecular dynamics simulations. The Journal of Chemical Physics 1995, 102, 5451–5459.

[60] Wassenaar, T. A.; Ingólfsson, H. I.; Böckmann, R. A.; Tieleman, D. P.; Marrink, S. J. Computational lipidomics with insane: A versatile tool for generating custom membranes for molecular simulations. Journal of Chemical Theory and Computation 2015, 11, 2144–2155.

[61] Šali, A.; Blundell, T. L. Comparative protein modelling by satisfaction of spatial restraints. Journal of Molecular Biology 1993, 234, 779–815.

[62] Humphrey, W.; Dalke, A.; Schulten, K. VMD: Visual molecular dynamics. Journal of Molecular Graphics 1996, 14, 33–38.

[63] Michaud-Agrawal, N.; Denning, E. J.; Woolf, T. B.; Beckstein, O. MDAnalysis: A toolkit for the analysis of molecular dynamics simulations. Journal of Computational Chemistry 2011, 32, 2319–2327.

[64] Pettersen, E. F.; Goddard, T. D.; Huang, C. C.; Meng, E. C.; Couch, G. S.; Croll, T. I.; Morris, J. H.; Ferrin, T. E. UCSF ChimeraX: Structure visualization for researchers, educators, and developers. Protein Science 2021, 30, 70–82.

