## Supplementary Information for "*Switching Gō-Martini* for Investigating Protein Conformational Transitions and Associated Protein-Lipid Interactions"

### 1 Supplementary Tables

**Table S1:** MD parameters for all CG simulation systems.

| Parameters | GBP | AdK | β2AR | ATP8A1-CDC50a |
|---|---|---|---|---|
| cutoff-scheme | verlet | | | |
| coulombtype | reaction-field | | | |
| epsilon_r | 15 | | | |
| rcoulomb (nm) | 1.1 | | | |
| vdw_type | cutoff | | | |
| rvdw (nm) | 1.1 | | | |
| tcouple | V-rescale | | | |
| tau_t (ps) | 1.0 | | | |
| ref_t (K) | 310.15 | | | |
| Pcoupl | Parrinello-Rahman | | | |
| Pcoupletype | isotropic | | semiisotropic | |
| ref_p (bar) | 1.0 | | 1.0, 1.0 | |
| tau_p (ps) | 12.0 | | | |
| | Minimization | | | |
| integrator | steep | | | |
| nsteps | 5000 | 5000 | 15000 | 15000 |
| | Equilibration | | | |
| intergrator | md | | | |
| dt (ps) | 0.020 | | | |
| nsteps | 5e4 | 5e4 | 1e5 | 5e5 |
| simulation time (ns) | 1 | 1 | 2 | 10 |
| | Production | | | |
| integrator | md | | | |
| dt (ps) | 0.020 | | | |
| nsteps | 1.5e7 * 3 | 5e6 * 2 | 2.5e7 * 2 | 1e7, 4e7 |
| simulation time (ns) | 300 * 3 | 100 * 2 | 500 * 2 | 200, 800 |
| repeats | 50 | 70 | 40 | 50 |
| | Transition relaxation | | | |
| integrator | md | | | |
| dt (ps) | 0.002 | | | |
| nsteps | 1e3 | | | |

**Table S2:** Transition pathway summary for all CG simulation systems.

| Protein | Pathway | Number of simulations |
|---|---|---|
| **GlnBP** | Open-closed pathway | 50 (100%) |
| **AdK** | NMP-closing pathway | 48 (68.6%) |
| | LID-closing pathway | 7 (10.0%) |
| | Middle | 15 (21.4%) |
| **β2AR** | Normal | 39 (97.5%) |
| | Abnormal | 1 (2.5%) |
| **ATP8A1-CDC50a** | Main | 36 (72%) |
| | Middle | 5 (10%) |
| | Incomplete | 9 (18%) |

### 2 Supplementary Figures

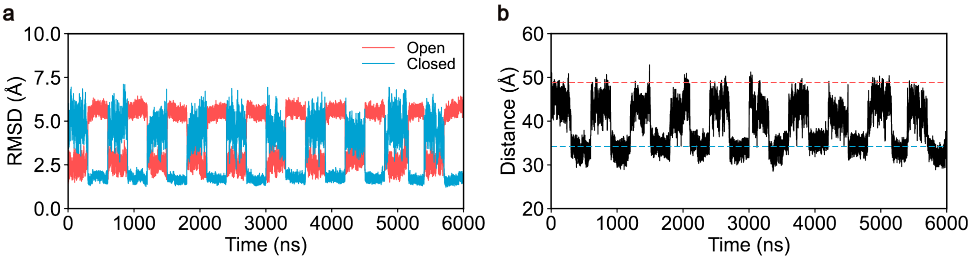


**Figure S1: Conformational transitions of GlnBP. a,** The RMSD of GlnBP during one simulation comprising 10 cycles of open⟷closed transitions, with respect to the open- (red) or closed-state (blue) structures. **b,** The distance between the backbone atoms of T59 and T130 during the simulation comprising 10 cycles of open⟷closed transitions. The red and blue dashed lines represent the reference distance from the open- and closed-state structures.

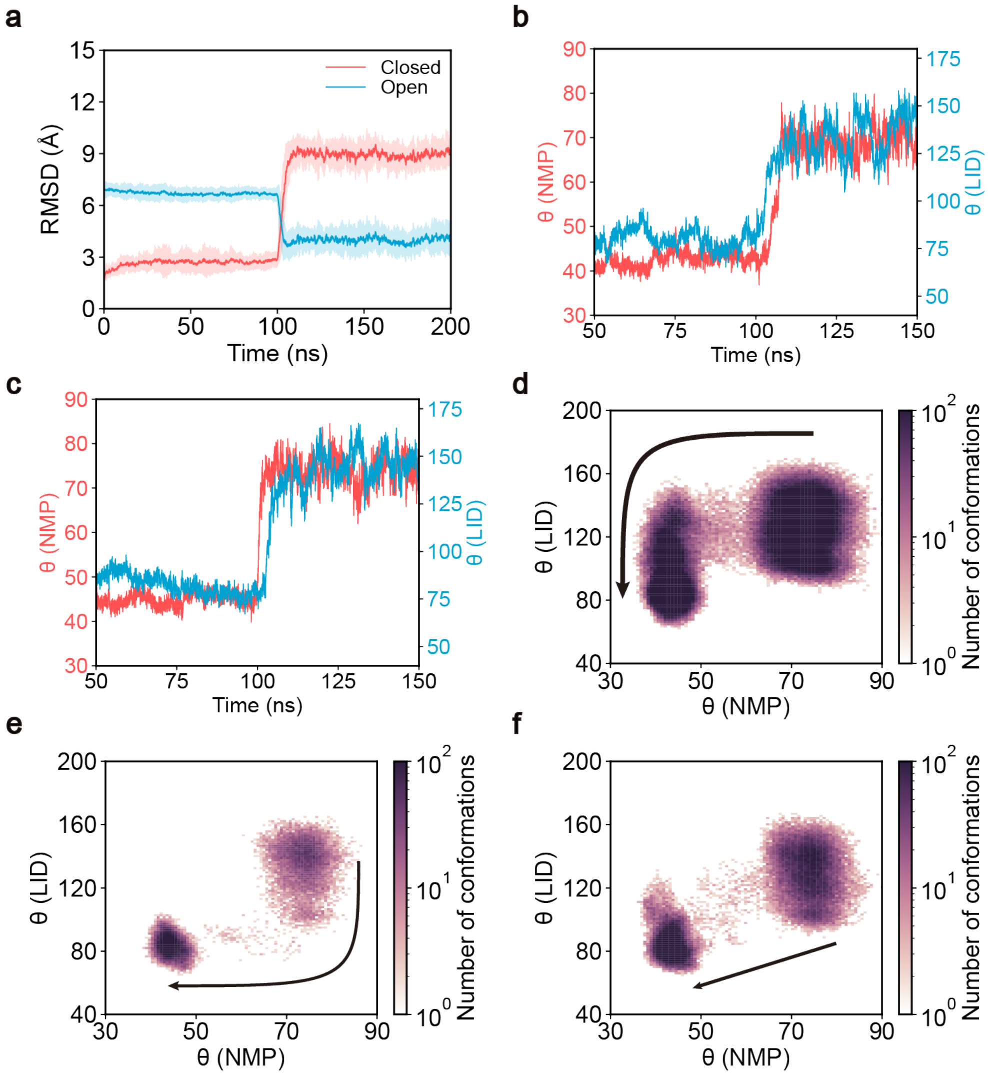


**Figure S2: Conformational transitions of AdK. a,** The RMSD of AdK with respect to the closed-state (red) and open-state (blue) structures along the simulations. The solid lines correspond to the average values obtained from 70 independent simulation trajectories, while the shaded area represents the standard deviation. **b,** Time series of $\theta_{NMP}$ (red, left y-axis) and $\theta_{LID}$ (blue, right y-axis) for AdK's NMP-closing pathway. The data used is the same as that in Fig. 2e. **c,** Time series of $\theta_{NMP}$ (red, left y-axis) and $\theta_{LID}$ (blue, right y-axis) for AdK's LID-closing pathway. The data used is the same as that in Fig. 2f. **d–f,** Two-dimensional histograms of AdK conformations using two angles $\theta_{NMP}$ and $\theta_{LID}$ as the reaction coordinates. The NMP-closing pathway, LID-closing pathway, and Middle pathway are shown in **d**, **e**, and **f**, respectively. The color represents the number of conformations sampled during all simulations for the respective pathways, with the max number cutoff of 100. The black curves represent simplified conformational transition flux, with the width representing the relative occupancy of the transition pathways.

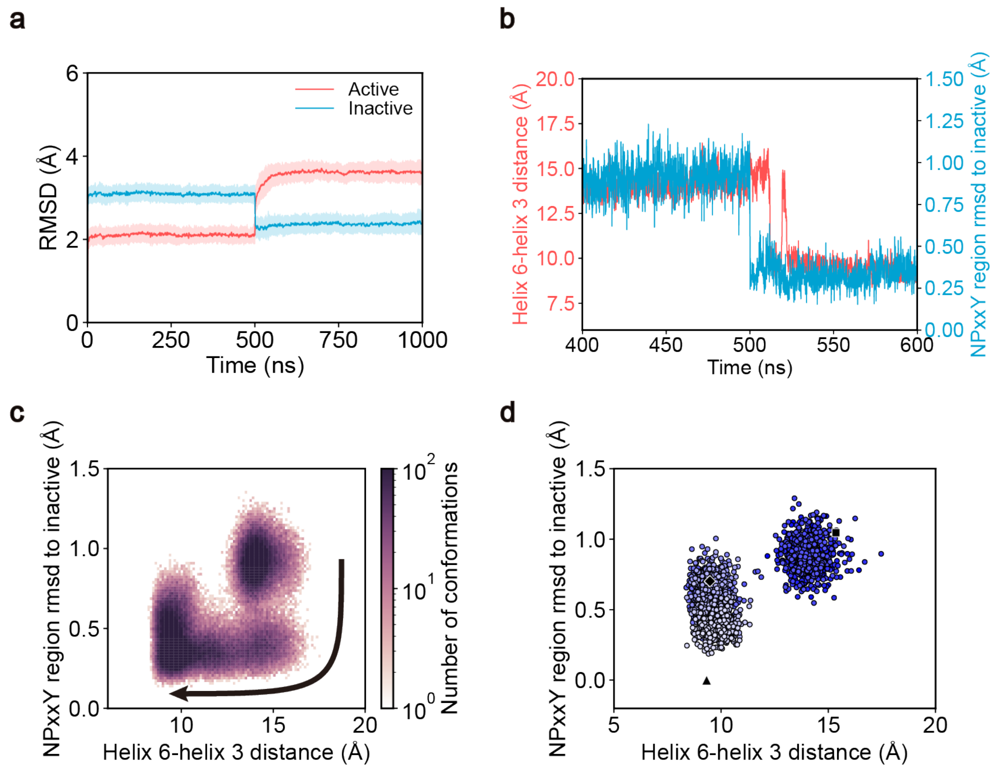


**Figure S3: Conformational transitions of β2AR. a,** The RMSD of β2AR during the simulations. The red and blue lines are RMSDs with respect to the active- and inactive-state structures of β2AR. The solid lines correspond to the average values obtained from 40 independent simulation trajectories, while the shaded area represents the standard deviation. **b,** Time series of the helix 6–helix 3 distance (red, left y-axis) and the RMSD of NPxxY motif with respect to the inactive state (blue, right y-axis) for β2AR's normal pathway. The data used is the same as that in Fig. 3c. **c,** The two-dimensional histogram of β2AR conformations of the normal pathway. The color represents the number of conformations sampled during all 39 normal pathway simulations, with the max number cutoff of 100. The black curve is the simplified conformational transition flux. **d,** The abnormal conformational transition pathway of β2AR, which is constructed with the helix 6–helix 3 distance and the RMSD of NPxxY motif with respect to the inactive state. The blue color of the circles decreases along with the simulation time. The black square, diamond, and triangle are representative of the active, intermediate, and inactive states of β2AR.

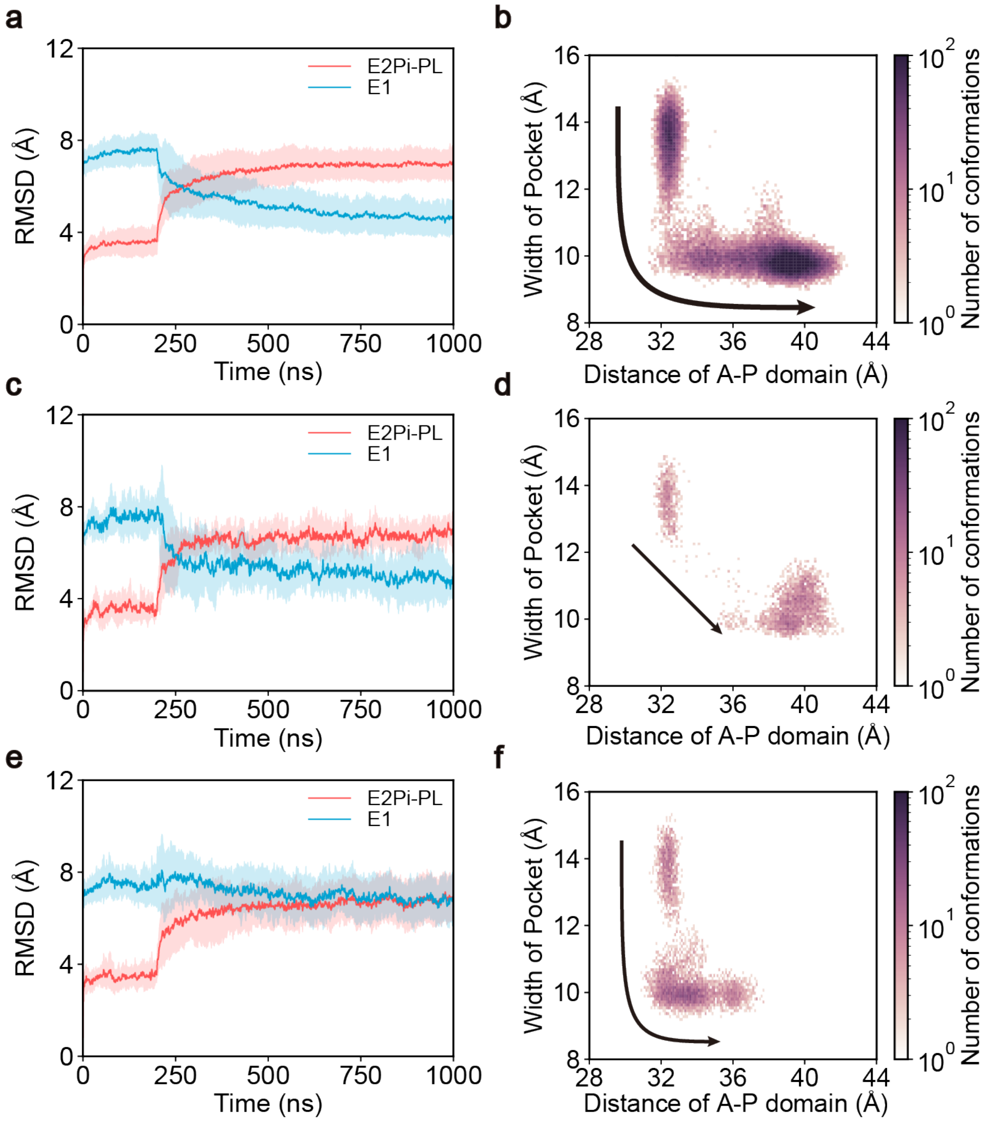


**Figure S4: Conformational transitions of ATP8A1-CDC50a. a,** The RMSD of ATP8A1-CDC50a during conformational transitions of the main pathway. The red and blue lines are RMSDs with respect to the E2Pi-PL and E1 state structures of ATP8A1-CDC50a. The solid lines correspond to the average values obtained from all respective simulation trajectories, while the shaded area represents the standard deviation. **b,** The two-dimensional histogram of ATP8A1-CDC50a conformations of the main pathway. The color represents the number of conformations sampled during all simulations for the respective pathway. The black curves represent simplified conformational transition flux, with the width representing the relative occupancy of the transition pathways. **c–f** are similar to **a, b**, but for the middle pathway and the incomplete simulations, respectively.

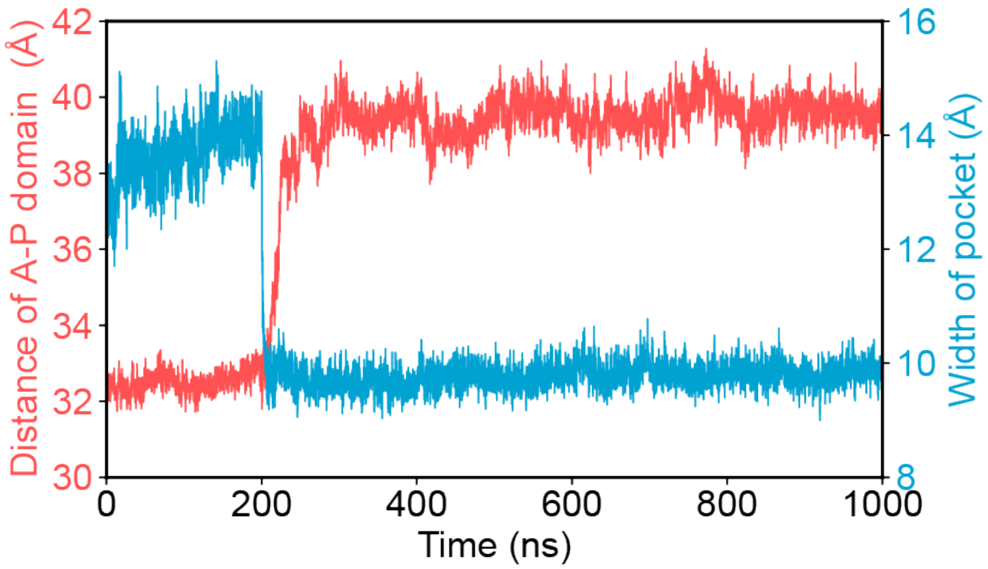


**Figure S5: Time series of the main pathway for ATP8A1-CDC50a.** Time series of the distance of A-P domain (red, left y-axis) and the width of the lipid-binding pocket (blue, right y-axis) for ATP8A1-CDC50a's main pathway. The data used is the same as that in Fig. 4c.

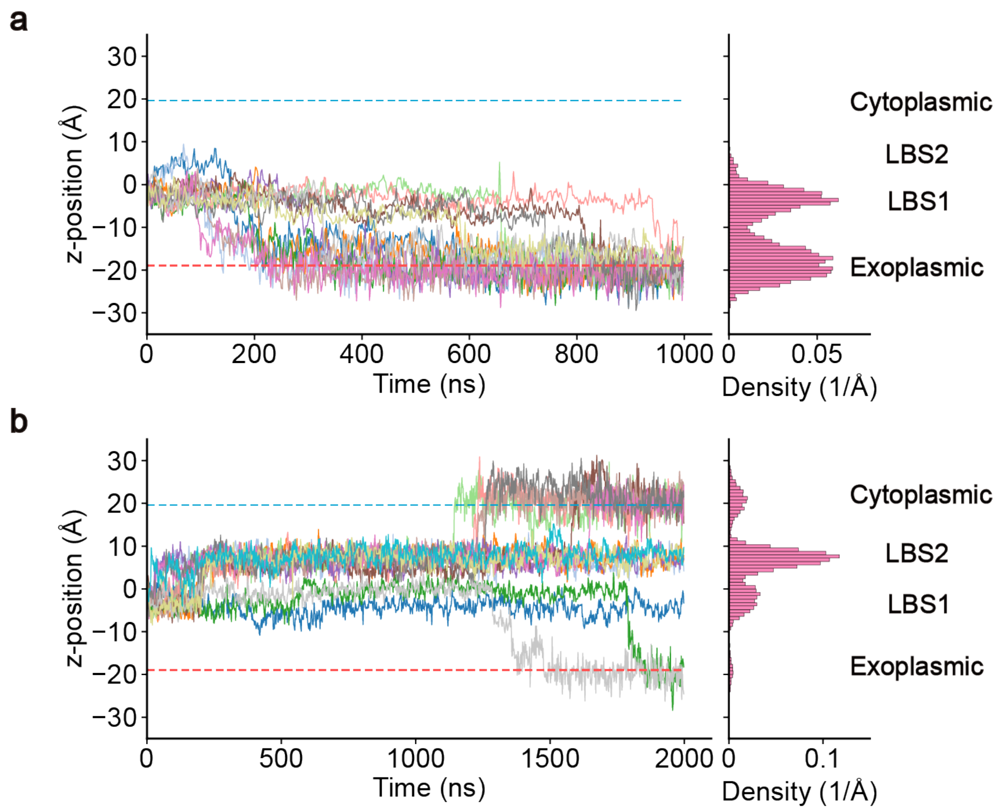


**Figure S6: Lipid translocation of ATP8A1-CDC50a. a, b,** Traces (left) and distribution density (right) of the lipid heads across the z-axis in the course of simulations. The failed and the incomplete lipid translocation for the first 1000 ns simulations are shown in **a** and **b**, respectively. For the incomplete lipid translocation, an additional 1000 ns simulation was conducted to investigate both the stability of the lipid trapping state and the flipping tendency for lipid translocation. The solid lines with different colors represent different simulations. Red and blue dashed lines represent the exoplasmic and cytoplasmic leaflets of the bilayer membrane. The positions of lipid binding sites (LBS1, LBS2) are also labeled.

#### 3 Supplementary Movie

**Movie S1. POPS lipid translocation process of ATP8A1-CDC50a**. The movie shows a representative trajectory of the POPS lipid translocation process by ATP8A1-CDC50a. Each domain of proteins is illustrated with the same color code as in Fig. 4. The POPS tail is colored medium orchid, and the lipid head group is shown as orange and red beads.
